# Resident soil microbial diversity and urea amendment legacy interact to shape the composition and expression of a surface film–forming soil inoculant

**DOI:** 10.64898/2026.06.08.730853

**Authors:** Ryan V. Trexler, Mary Ann Bruns, Mikayla A. Borton, Jason P. Kaye, Estelle Couradeau, Terrence H. Bell

## Abstract

Soil microbial inoculants have the potential to improve crop yield, enhance agricultural sustainability, and support soil restoration, but they often display unpredictable in-field performance across varied soil conditions. Cyanobacteria-dominated soil surface consortia (SSCs) offer a tractable model for studying inoculant-soil interactions because their visible surface growth enables direct observation and sampling after application. Here, we introduced the SSC “DG1,” dominated by the diazotrophic cyanobacterium *Nostoc linckia*, into soil microcosms differing in resident microbiome diversity (low vs. high diversity) and urea fertilization history, (+urea vs. –urea). We used 16S rRNA gene sequencing and genome-resolved metatranscriptomics to assess inoculant establishment and functioning. Resident microbiome diversity did not affect total *N. linckia* gene expression, but heterotrophic DG1 members showed reduced expression in high-diversity soils. Soil diversity and urea history drove broad shifts in DG1 transcription and significantly affected transcription of key *N. linckia* carbon and nitrogen metabolism genes. High-diversity soils with urea were associated with increased transcription of photosynthesis, CAZyme, and nitrogen cycling genes, whereas low-diversity soils without urea promoted increased nitrogenase transcription and reduced carbon and nitrogen metabolism transcription. These results show that inoculant outcomes depend not only on establishment, but also how soil conditions and biology shape post-establishment functioning.

## Introduction

Microbial inoculants are living microorganisms that have been captured in one environment and introduced to another, often at high concentrations, to produce desirable effects in the recipient environments. In agriculture, this approach has been used to augment soil functions for more than 100 years^1^, but interest by the agricultural biotechnology industry has surged in the past decade. This shift is coincident with innovations in high-throughput sequencing and microbial therapeutics in human health, and is driven by goals of improving crop yields, advancing sustainable agriculture, and supporting soil restoration efforts^2,3^. Many commonly sold inoculants are difficult to observe after their application, making it challenging to track their survival and performance without molecular tools, but some emerging inoculants do not share this limitation. Naturally-occurring cryptobiotic soil covers from agroecosystems, as well as dryland environments^4^, are being investigated for use in sustainable agriculture^5,6^ and dryland ecological restoration^4^, respectively.

In agroecosystems, such soil covers are formed by assemblages of organisms that form ephemeral biofilms on soil surfaces and are generally dominated by photosynthetic Cyanobacteria. Cyanobacterial-based inoculants have been explored in regions such as South Asia^7^, but such covers are also widespread in temperate agroecosystems. Recent efforts^5,6,8^ have used enrichments in nitrogen- and carbon-free media to capture various consortia, referred to as soil surface consortia (SSC), in the Northeast United States. One particular SSC, Dark Green 1 (DG1), demonstrated high photosynthetic efficiency and was shown to modulate soil nitrogen availability^6^ and stabilize soil surfaces^5^. Within its enrichment culture, DG1 is dominated by the diazotrophic cyanobacterium *Nostoc linckia* (∼85% relative abundance) and includes six other heterotrophic bacteria (see results). Because of its ability to fix carbon and nitrogen, role in soil stabilization, and natural occurrence, DG1 is a promising soil surface inoculant for temperate agriculture systems.

Though the use of soil inoculants holds promise for the management of in-field soil microbiomes, unpredictable product performance in wild soils has limited reliance on inoculants by land managers. This unpredictability is thought to be due, in part, to differences between the abiotic and biotic conditions of the environment of origin and those of recipient environments^9^. The processes of isolating and producing inoculant biomass in nutrient rich monocultures may also reduce the success of establishment and function in new soil environments^10^. At present, a primary barrier to developing widely predictable and effective inoculants is a lack of understanding about how interactions between inoculants, established soil organisms, and soil physicochemical conditions affects inoculant establishment and function.

Due to their abundant and easy to visualize surface growth, SSC inoculants are unique in that their establishment and persistence can easily be tracked and visually sampled at will. Thus, SSCs represent a tractable and important model system for examining inoculant interactions with soil conditions. In this study, we introduced the SSC DG1, developed for use in temperate soils, to soil microcosms differing in the presence or absence of a diverse resident soil microbiome (low vs. high diversity) and in nitrogen status (+ urea vs. -urea), resulting in a 2×2 factorial design. After a period of establishment, the DG1 consortium was sampled to determine its in-soil composition (16S rRNA gene sequencing) and in-soil function (metatranscriptomic sequencing). Using metagenomes of the initial DG1 enrichment, we used a MAG-resolved metatranscriptomics approach to determine how variation in resident soil microbial diversity and nitrogen fertilization history influences inoculant establishment and functioning. We hypothesized that resident soil microbial diversity and nitrogen fertilization history will independently and interactively effect 1) DG1 establishment on soil microcosms, and 2) DG1 gene transcription, particularly those related to carbon and nitrogen cycling. Uniquely, we are able to demonstrate that the resident soil microbiome and urea fertilization history interact to shape both the establishment and gene transcription of the DG1 inoculant and the expression of key carbon and nitrogen metabolism genes by the DG1 inoculant cyanobacteria *N. linckia*. This suggests that soil physicochemical and biological conditions jointly govern the establishment and functioning of soil inoculants, indicating that predictions of in-field performance must account for both soil properties and the resident microbial community.

## Materials and Methods

### Microcosm establishment and harvesting

A microbiology resource announcement describing the experimental design and establishment of soil microcosms has been published^11^. Briefly, we added DG1 to soil microcosms containing Low or High Diversity soil microbiomes, with or without soil amended with urea (Supplemental Figure 1, panel A). This resulted in a 2×2 factorial design with 4 treatments: 1) Low diversity soil, no added urea; 2) low diversity soil, +urea; 3) high diversity soil, no added urea; 4) high diversity soil, +urea. A total of 6 replicate microcosms were prepared for each treatment type. The soil used for preparing microcosms was collected from the Pennsylvania State Research Agronomy Farm at Rock Springs, which has been characterized previously^5^, and is classified as Hagerstown silt-loam. The soil was dry sieved to 2 mm, and twice autoclaved (45 min, 20 psi, 120 °C) with a 24-hour resting period between cycles. We have shown that autoclaving is as effective as gamma-irradiation for microbial recolonization experiments^12^; although both approaches impact soil physicochemistry, this provides a much more realistic model substrate than culture media. To eliminate differences in soil chemistry due to autoclaving^12^ across treatments, all microcosms were prepared from the same batch of autoclaved soil. The autoclaved soil was used directly for the Low Diversity microcosms (i.e. only organisms that survived autoclaving and grew back would be present), while soil for the High Diversity microcosms was prepared by re-inoculating the autoclaved soil with fresh, non-autoclaved soil at a ratio of 5% (volume fresh soil / volume autoclaved soil). From the Low and High Diversity soils, 25 g of dry soil was distributed to 12 Petri dishes (10 x 15 mm), resulting in 24 total microcosms. An even mixture of fructose/maltose/glucose/galactose/ribose (calculated to provide even carbon mass from each substrate) was added to all microcosms at a rate of 2 g carbon/kg dry soil. To establish treatments with added urea, 6 of the 12 microcosms of each soil type also received urea at a rate of 50 mg N/kg dry soil. All 24 microcosms were given a substantial amount of time to acclimate post-treatment and were incubated in the dark at 21°C for a total of 43 weeks to establish relatively stable soil microbiomes.Each microcosm was non-destructively sampled for DNA extractions (see below) after 43 weeks of incubation in order to obtain baseline soil microbiome data prior to DG1 addition to the microcosms.

As described previously^5^, the DG1 consortium used in this study was enriched from a naturally-occurring agricultural SSC that was collected from the Penn State Agronomy Research Farm at Rock Springs, PA, USA. DG1 was enriched in modified, carbon- and nitrogen-free BG-11 media over 2 years. Prior work on DG1 reported that the consortium was dominated by one or more strains of *Cylindrospermum* spp. and at least 6 other non-photosynthetic bacteria^5^ based on the alignment of 16S rRNA genes from metagenome-derived bins. Here, however, MAG-level ANI-based taxonomy reveals the dominant Cyanobacteria member is closely related to *Nostoc linckia* (see Results for more details).

A starter culture of the DG1 consortium was initially grown from frozen stock in 25 mL of carbon- and nitrogen-free modified BG-11 media. We then generated sufficient DG1 biomass for use in soil microcosms by transferring 2.5 mL of the starter DG1 enrichment culture into 250 mL of carbon and nitrogen free modified BG-11 media. All cultures of DG1 enrichment were incubated for 4 weeks with moderate agitation in baffled flasks at 21 °C and under fluorescent lighting with an average of 1,865 lux. The cultures were transferred to 50 mL conical vials and centrifuged at 5500 rpm to pellet the biomass. The supernatant was removed and the pellet was resuspended with sterile deionized water (3:1 vol/vol). We pipetted 3 mL of the resuspended biomass across the surface of the soil in each established microcosm and incubated the microcosms at 21°C for 5 weeks under fluorescent light at an average of 1,865 lux. Microcosms were sealed with parafilm to minimize water loss.

The microcosms were destructively harvested after 5 weeks of additional incubation with the DG1 consortium on the soil surface. We added 1 mL sterile deionized water to each microcosm 8 days before harvesting to promote active transcription by soil microorganisms and DG1 consortia members. For each of the 24 microcosms, a portion of the DG1 soil surface film, as well as directly embedded soil aggregates, was excised using a sterile razor blade and cut into two subsamples, each placed in a microcentrifuge tube (Supplemental Figure 1, panel B-C).

### DNA extractions and 16S rRNA gene library preparation

DNA extractions were completed on samples from both the microcosm soil prior to DG1 addition and the excised soil surface film (Supplemental Figure 1). DNA extractions, 16S rRNA gene amplicon library preparation, and Illumina sequencing was performed as described previously^13^. Specifically, DNA was extracted using the NucleoSpin soil 96 kit (Macherey-Nagel) using ∼300 mg of the excised soil film and embedded soil aggregates. Lysis disruption was performed for 30 s at 4.0 m/s using the FastPrep24 homogenizer (MP Biomedicals) with Buffer SL and without Enhancer SX. The v4 region of the 16S rRNA gene was PCR amplified using universal bacterial primers^14,15^ 515F (5’-GTGYCAGCMGCCGCGGTAA-3’) and 806R (5’-GGACTACNVGGGTWTCTAAT-3’) that were designed with overhangs to attach barcodes and standard Illumina overhang adaptors during a second PCR step^16^. Barcoded amplicons from the second PCR step were normalized using the SequalPrep Normalization plate kit (Invitrogen, Carlsbad, CA) and subsequently pooled. The pooled barcoded PCR product was then concentrated using a Savant SpeedVac at 50°C for 3 h. The concentrated pool was separated on a 1.2% agarose gel and the expected band excised from the gel before being purified with the PureLink Quick Gel extraction kit (Invitrogen). Illumina sequencing of the 16S rRNA gene amplicon libraries was done at the Cornell University Biotechnology Resource Center Genomics Facility using an Illumina MiSeq and 2 x 250 cycle v2 kit. Raw 16S rRNA amplicon reads are available through the NCBI Short Read Archive under the project number PRJNA547780.

### 16S rRNA gene ASV analysis

A total of 2,205,725 raw paired-end reads were processed for determining 16S rRNA gene Amplicon Sequence Variants (ASVs) for samples, both before and after DG1 addition (Supplemental Figure 1). First, cutadapt^17^ was used to trim Illumina sequencing adapters and the DADA2^18^ pipeline was implemented in R to generate ASVs. DADA2 was used to filter chimeric sequences and assign taxonomy using the Greengenes 13.8 database^19^. The R *vegan* package^20^ was used to determine diversity indices, calculate Bray-Curtis distances from the ASV table, and construct non-metric multidimensional scaling (NMDS) ordination. The effect of treatment on the dissimilarity of community composition (Bray-Curtis ∼ Soil Diversity * Nutrient Status) was tested by PERMANOVA^21^ via the *vegan::adonis2* function. Bacterial composition was assessed using the *phyloseq* package^22^ and measures of ASV richness and Shannon diversity were tested by ANOVA followed by Tukey’s post hoc test to compare treatments. Data was visualized using the *ggplot2* package^23^. In order to assign 16S rRNA gene ASVs to DG1 MAGs, ASV sequences were aligned (>98% identity) via BLAST^24^ to MAG contigs containing 16S rRNA genes and by matching ASV sequence taxonomy to MAG taxonomy.

### DG1 consortium metagenome sequencing, MAG curation, and MAG annotation

Metagenomic sequencing from 3 technical replicates of the DG1 consortium was performed in Peng and Bruns 2019^5^ using the Illumina MiSeq and 150bp v2 kit and submitted to GenBank under accession SAMN04488150. In this study, raw metagenome reads were quality trimmed using sickle v1.33^25^ and the forward and reverse reads of the 3 technical replicates were concatenated to a fasta file for the forward and reverse reads. The combined technical replicates were co-assembled via metaSPAdes v3.15.5^26^ and the subsequent contigs were binned using metaBAT2 v2.17^27^. CheckM2 v1.0.2 was used to assess bin quality according to the minimum information about metagenome-assembled genomes (MIMAG) standards^28^ and all 7 bins were retained in the resulting metagenome-assembled genome (MAG) database. Missing rRNA genes for each MAG were attempted to be retrieved by aligning the coassembly with a database of MAG-related rRNA genes via BLAST^24^ and then by manual curation of the identified contigs. The quality filtered reads were mapped with *bbmap.sh*^29^ to the MAG database in order to assess the amount of the metagenome captured by the MAG database.

MAGs were annotated with DRAM v2^30^, where open reading frames (ORFs) were identified by Prodigal^31^ and ORFs were searched against the KEGG^32^, UniRef90^33^, MEROPS^34^, Pfam^35^, dbCAN^36^, VOGdb^37^, KOfam^38^ databases using MMSeqs2^39^ and HMMER^40^. Additionally, DRAMv2 uses tRNAscan-SE^41^ and barrnap (https://github.com/ tseemann/barrnap) to identify tRNA and rRNA genes, respectively. The taxonomy of each MAG was identified via GTDB-Tk v2.4.0^42^ and GTDB release220. Lastly, singleM v0.18.3^43^ was used to estimate the relative abundance of each DG1 MAG in the three metagenome replicates. Here, the singleM pipe command was run on the trimmed forward and reverse metagenome reads to generate taxon tables for each replicate. The singleM summarise command was run to combine the replicate taxon tables, which were subsequently analyzed in R using phyloseq^22^, in which representative sequences from singleM were matched to each MAG via taxonomy.

### Metatranscriptome library preparation and sequencing

RNA extractions were performed on the excised soil surface film immediately after harvesting using the RNeasy PowerSoil total RNA (Qiagen) kit, and the quality of the extracted RNA was evaluated using the Agilent BioAnalyzer at the Penn State University Genomics Core (RNA Integrity Number [RIN] > 7). RNA samples were shipped on dry ice to the United States Department of Energy (US-DOE) Joint Genome Institute (JGI) for metatranscriptome library preparation, which was described previously^11^. Briefly, rRNA was depleted using the Ribo-Zero rRNA (Illumina) removal kit (equimolar bacteria, yeast, plant root) and libraries were constructed with the TruSeq stranded total RNA high-throughput kit using 100 ng/sample RNA, and Polymerase Chain Reaction (PCR) amplified with 10 cycles. The Roche LightCycler 480 and KAPA library quantification kit were used to quantify the prepared libraries. Sequencing was performed on the Illumina NovaSeq with the XP v.1 reagent kit allowing for an indexed 2 x 150 bp cycle run. Raw reads are available via the Joint Genome Institute Gold database under GOLD study ID Gs0132857.

### Metatranscriptome processing, MAG-resolved transcription, and downstream analysis

A total of 5,154,599,632 raw paired-end reads were obtained across the 24 metatranscriptome samples. Reads were quality trimmed with sickle^25^ and adapter sequences and remaining low quality reads were removed with bbduck.sh^29^. Reads were further filtered by RQCfilter^29^ to remove potential contaminant sequences. In all, a total of 4,833,069,406 quality-assured reads were obtained for the 24 metatranscriptomes and subsequently mapped using bowtie2^44^ to the scaffolds of the 7 DG1 MAGs. SAMtools^45^ and bbmap::reformat.sh^29^ were used to format the resulting mapping files and featureCounts^46^ was used to count the number metatranscriptome reads mapped to each DG1 MAG gene. From the featureCounts mapping table, Reads per kilobase (RPK) values for each gene were calculated in R and then EdgeR^47^ was used to compute GeTMM^48^ expression values for each gene following a previously published pipeline^49^.

For each of the seven DG1 MAGs metagenome-derived DRAM annotation data was merged to the GeTMM expression data from the 24 metatranscriptome samples in R. The DRAM v2 genome_summary_form.csv was used to perform custom distillation and summarization of the annotated metatranscriptome data. Further, the *vegan* package^20^ was used to Hellinger-transform the GeTMM expression data and compute a Redundancy Analysis (RDA) to constrain multivariate ordination space by Soil Resident Diversity and Nutrient Amendment (GeTMM.hellinger ∼ Soil Resident Diversity * Nutrient Amendment). The effects of Soil Resident Diversity and Nutrient Status on DG1 MAG expression patterns was tested via *adonis2* in the *vegan* package^20^ (GeTMM.hellinger ∼ Soil Resident Diversity * Nutrient Amendment).

The expression of the *Nostoc linckia* MAG was of particular interest, given its key role within the consortium, so we performed a targeted analysis of transcripts related to photosynthesis, carbon-fixation, nitrogen fixation, heterocyst formation, nitrogen metabolism, CAZymes, and extracellular polysaccharide production. Gene lists for these functions (Supplemental file 1) were compiled from both the DRAM2 genome_summary_form.csv and from manual curation based on the literature. The gene lists were used to manually pull matching genes from the *Nostoc linckia* MAG DRAM2 annotation. Heatmaps were built using the ComplexHeatmaps package^50^ with z-scores calculated from GeTMM expression per gene across the 24 metatranscriptome samples. Analysis of differential expression of these genes was performed in R using the *limma* package^51^ via linear regressions (GeTMM ∼ Soil Resident Diversity * Nutrient Amendment). In the case of multiple copies of target genes, homologs were collapsed by summing the GeTMM expression for the gene within each sample before input into *limma*. Gene-wise sample variance was minimized through empirical Bayes moderation using the *limma eBayes()* function. The same methodology was used to run the differential expression analysis at the pathway level where GeTMM values were summed across pathway genes. Linear regressions (GeTMM_pathway_ ∼ Soil Resident Diversity * Nutrient Amendment) were again computed via the *limma* package in R. Additionally, the *limma makeContrasts()* function was used to perform pairwise contrasts between all combinations of treatments, and the resulting BH adjusted p-values were used to indicate significant groupings by treatment.

## Results

### Microcosm microbial diversity and composition

Following incubating the microcosm soils in the dark for 43 weeks, and prior to adding the DG1 inoculant to these microcosms (Soil Only Sampling Phase), we verified that the Low and High Diversity treatments harbored contrasting soil microbiomes (Figure 1A and Supplemental Figure 2). Here, we confirmed the Low Diversity treatments had significantly lower ASV richness and Shannon Diversity compared to High Diversity treatments. Within each of the two resident soil diversity treatment levels, alpha diversity did not significantly differ between Urea Amendments, with the exception of Simpson’s Evenness which was higher in the Low Diversity Urea (-) treatment compared to Low Diversity Urea (+) (Supplemental Table 1 and Supplemental Figure 2). The resident soil bacterial composition in the High Diversity treatments were marked by a diverse set of common soil taxa, however, the Low Diversity treatments were largely composed of only a few ASVs from the Actinobacteria and Firmicutes, though two microcosms in the Low Diversity Urea (+) treatment harbored ASVs from additional phyla (Supplemental Figure 3).

**Figure 1:**
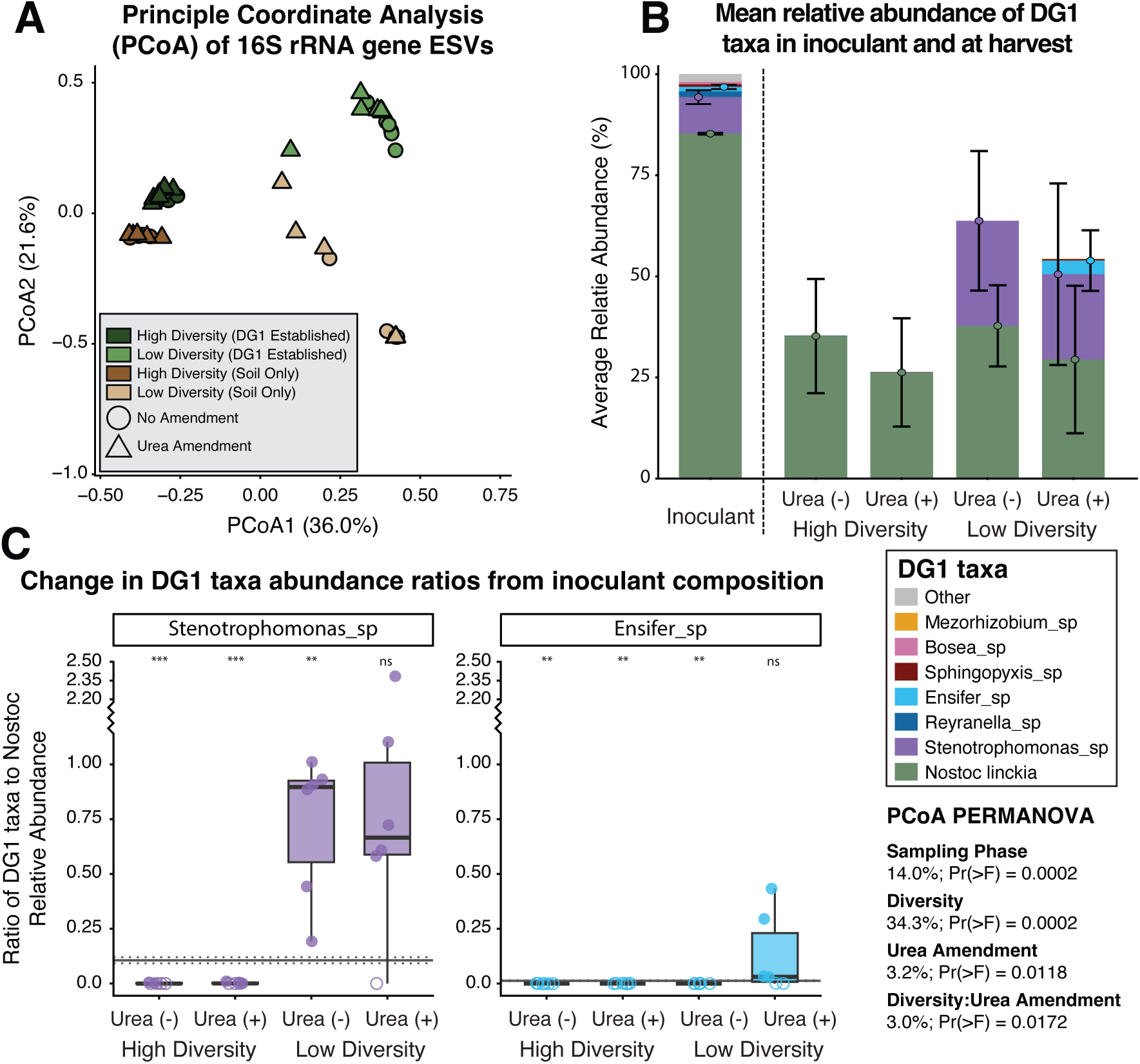
**A)** Principal Coordinates Analysis (PCoA) of 16S rRNA gene ASVs among all treatments using Bray-Curtis dissimilarities. Brown (High Soil Diversity) and tan (Low Soil Diversity) points indicate microcosms prior to DG1 addition, while microcosms at the end of the experiment with DG1 added are displayed in Dark Green (High Soil Diversity) and Light Green (Low Soil Diversity). Triangles represent microcosms that received urea amendment, while circles show microcosms with no added urea. A PERMANOVA (bottom right) was performed across all samples, demonstrating significant effect of Sampling Phase, Soil Diversity, Urea Amendment, and Diversity:Urea Amendment on microbiome composition (α=0.05). **B)** Average relative abundance of DG1 MAGs in the inoculant (based on singleM) and across treatments at harvest (via 16S rRNA ASVs). Points with error bars indicate treatment means per MAG **±** 1 standard deviation for *N. linckia*, *Stenotrophomonas* sp., and *Ensifer* sp. **C)** Ratios of the relative abundance of each MAG to the relative abundance of *N. linckia* at the time of final harvest (points and boxplots). The solid line indicates the baseline mean ratio of the relative abundance of each MAG to the relative abundance of *N. linckia* in the starting DG1 inoculant. Dashed lines above and below the solid line shows the baseline mean ratio **±** 1 standard deviation. Points displayed as filled circles indicate the MAG was detected in the microcosm, while points displayed as unfilled circles indicate the MAG was not detected in the microcosm. The results of Welch t-tests with Empirical Bayes moderated baselines are summarize above each treatment, where *** = p < 0.001, ** = p < 0.01, * = p < 0.05, and ns = p > 0.05.

After DG1 inoculation and five additional weeks of incubation under light, green surface films formed by the DG1 inoculant were visible on all microcosms. At this stage (DG1 Established), the bacterial community composition of microcosms shifted (Figure 1A, PERMANOVA: Pr(>F) = 0.0002, R^2^ = 14.0%) compared to their composition prior to DG1 inoculation (Soil Only), largely due to the establishment of DG1 taxa (Supplemental Figure 3). The bacterial community composition of microcosms varied mainly by the Soil Diversity treatment (Pr(>F) = 0.0002, R^2^ = 34.3%), while Urea Amendment (Pr(>F) = 0.0118, R^2^ = 3.2%) and the interaction between Soil Diversity and Urea Amendment (Pr(>F) = 0.0172, R^2^ = 3.0%) each explained a smaller proportion of variation. Trends in alpha diversity following DG1 establishment remained similar compared to before DG1 was inoculated (Supplemental Figure 2), with microcosms of each Soil Diversity treatment (High/Low) continuing to harbor contrasting resident soil diversities. Further, Cyanobacteria ASVs were abundant in all microcosms at the final sampling (>15% relative abundance).

### DG1 metagenome assembled genomes

In total, we generated three high quality MAGs, three medium quality MAGs, and one low quality MAG from the three DG1 metagenome technical replicates according to MIMAGs standards^28^. Bins from these metagenomes have been previously published^5^, though, here we report an improved set of MAGs from these metagenomes with all but one MAG (*Mesorhizobium* sp.) >94% estimated completion and < 2% contamination (Supplemental Table 1). The DG1 inoculant is largely dominated by the cyanobacterium *Nostoc linckia* (85% relative abundance) and, to a lesser degree, a MAG classified as *Stenotrophomonas* sp. (9% relative abundance). The remaining MAGs and “Other” each account for less than 2% of the total bacterial relative abundance (Supplemental Figure 4 and Supplemental Table 2).

### Relative abundance of DG1 members across treatments

To determine the relative abundance of DG1 members in sampled microcosms, 16S rRNA gene ASVs were matched to DG1 MAGs by blasting ASV sequences to the DG1 MAGs and linking taxonomy. Among the seven DG1 MAGs, *N. linckia* was the only DG1 member detected in all the microcosms from both High Diversity treatments (+/- urea). Further, *N. linckia* reliably established in all microcosms regardless of underlying Soil Diversity or Urea Amendment. *Stenotrophomonas* sp. was present in Low Diversity treatments regardless of urea amendment, while *Ensifer* sp. was detected in 4/6 of the Low Diversity Urea (+) microcosms (Figure 1B-C). Additionally, compared to the starting inoculant, *Stenotrophomonas* sp. increased in relative abundance relative to *N. linckia* in the Low Diversity Urea (-) treatment (Figure 1C). Similarly, samples from 5/6 Low Diversity Urea (+) microcosms for *Stenotrophomonas* sp. and 4/6 for *Ensifer* sp. show increased relative abundance ratios to *N. linckia* compared to the starting inoculant; however, at the treatment level this result is insignificant, at least in part due to the failure of *Stenotrophomonas* sp. and *Ensifer* sp. to establish in 1 and 2 microcosms respectively. ASVs for all other DG1 members were infrequently detected in any of the treatments (Supplemental Figure 5), indicating failure of these members of the inoculant to establish.

### Patterns in transcription among DG1 members across treatments

The experimental treatments had a marked effect on the expression patterns of the DG1 organisms (Figure 2). As expected, considering its high relative abundance in the DG1 inoculant, *N. linckia* had by far the highest average gene transcription among the DG1 MAGs (Figure 2A). Although the average gene transcription of *N. linckia* was only affected by the Low Diversity Urea (-) treatment where it was lowest, the proportion of its transcribed genes was highest in this treatment (Figure 2B). *Stenotrophomonas* sp. had moderate mean gene transcription, and ∼50% of its genome transcribed, but only in the Low Diversity treatments regardless of the Urea amendment. Similarly, *Ensifer* sp. had mean gene transcription, but only in the Low Diversity Urea (+) treatment. The remaining MAGs (*Bosea* sp., *Sphyngopyxis* sp., *Reyranella* sp., and *Mesorhizobium* sp.) had a negligible total expression and proportion of their genomes transcribed across all treatments (Supplemental Figure 6). These transcription patterns track the 16S rRNA gene abundance data where *Stenotrophomonas* sp. and *Ensifer* sp. were not detected in High Diversity treatments and other MAGs were infrequently detected. Furthermore, RDA ordination demonstrated that a large portion of the total multivariate variation in DG1 MAG gene transcription was explained the experimental treatments (R² = 0.492; adj. R² = 0.415; permutation test: F(3,20)=6.45, p=0.001, 999 permutations), and PERMANOVA tests showed that the Soil Resident Diversity (R^2^ = 27.8%, P(>F) = 0.001) is the dominant factor explaining this variation, whereas Nutrient Status and the interaction of Soil Resident Diversity:Nutrient Status explained 12.2% (P(>F) = 0.001) and 9.5% (P(>F) = 0.001), respectively (Figure 2C).

**Figure 2:**
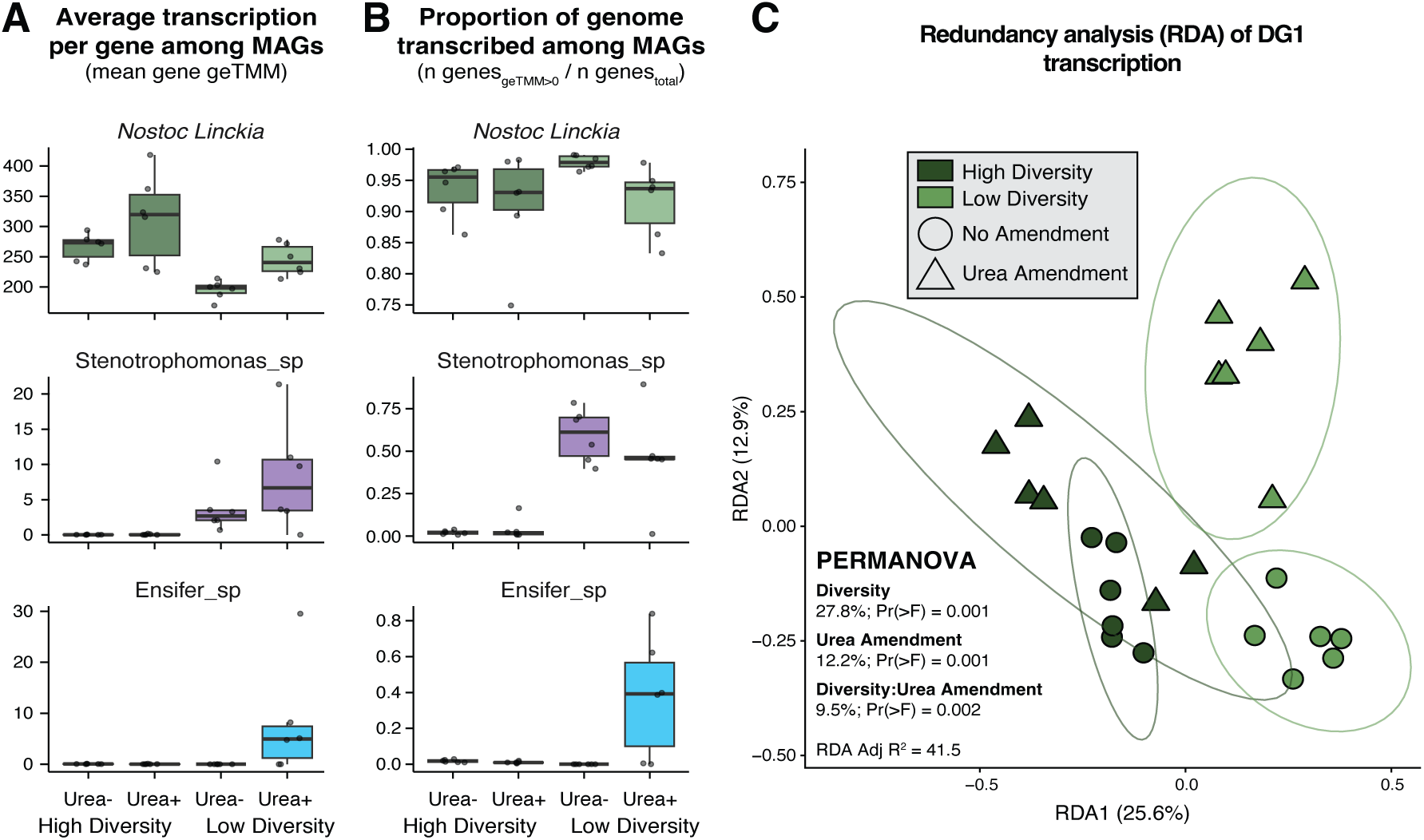
**A)** Boxplots showing the average gene geTMM transcription values for *N. linckia*, *Stenotrophomonas* sp., and *Ensifer* sp. between treatments. **B)** Boxplots showing the proportion of the genes per MAG that are transcribed for *N. linckia*, *Stenotrophomonas* sp., and *Ensifer* sp. between treatments. **C)** Redundancy analysis ordination of hellinger-transformed geTMM transcription values for all genes among the DG1 MAGs. Points for High Diversity samples are visualized in Dark Green and Low Diversity in Light Green. Urea-amended samples are marked by triangles and unamended samples by circles.

### Transcription of Nostoc linckia photosynthesis and carbon fixation genes

Considering the capacity of *N. linckia* to potentially add carbon to soil through photosynthesis, we explored the transcription of photosynthesis and carbon-fixing genes in *N. linckia,* specifically. Multiple linear regression models demonstrated that both Soil Diversity, Urea Amendment, and the interaction of both had significant effects (α=0.05) on the transcription of photosynthesis and carbon-fixing genes (Figure 3). At the pathway level, High Diversity soils promoted increased transcription of Photosystem I genes compared to Low Diversity soils and within Low Diversity microcosms, urea addition promoted an increased transcription of Photosystem I genes. High Diversity soils also showed higher transcription of Photosystem II, cytochrome b6f, and photorespiration genes than Low Diversity Urea (-) microcosms, but this difference was diminished under urea addition, as High Diversity Urea (+) and Low Diversity Urea (+) were similar for these pathways. High Diversity Urea (-) also had higher total transcription of RuBisCO and carboxysome genes compared to Low Diversity Urea (-), but under urea fertilization there was no difference between diversity treatments. Interestingly, no treatments led to a statistical increase in inorganic carbon import.

**Figure 3:**
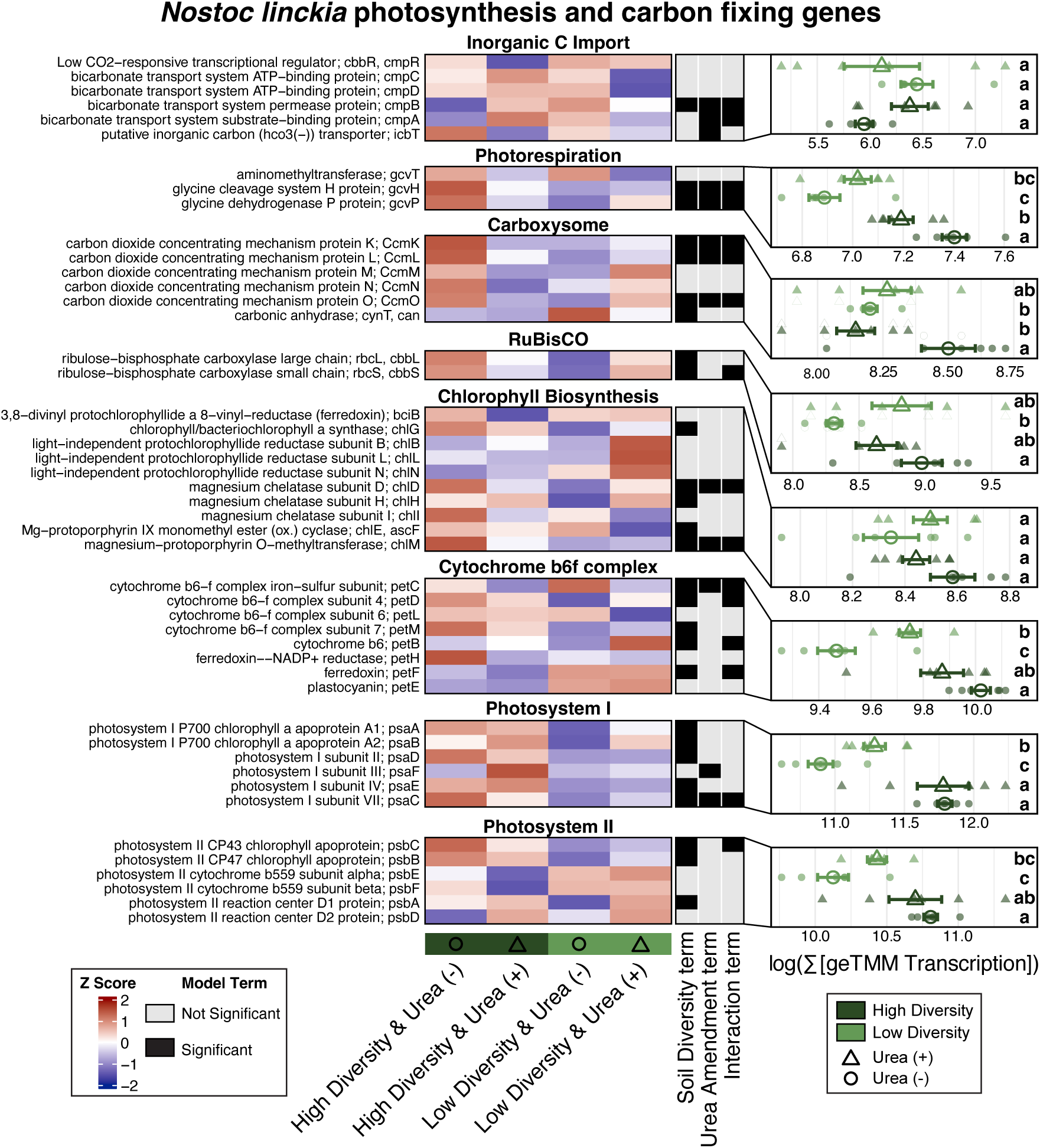
Heatmaps in left-most column represent z-scores of geTMM transcription values at the gene level (rows) for *N. linckia* photosynthesis and carbon fixing genes across the four treatments and faceted by pathways. Annotations at the gene level in middle column indicate significance (α=0.05, B-H corrected p-values) of model terms from multiple linear models across each gene. Right-most column shows the log of summed geTMM transcription values of genes among each pathway. Significance (α=0.05, B-H corrected p-values) between treatments were determined using ANOVA with Tukey-HSD post hoc tests and represented by grouping letters.

Additionally, we investigated the effect of Soil Diversity and Urea Amendment on the transcription of CAZyme genes in order to probe how *N. linckia* was potentially metabolizing organic carbon. Differential abundance analysis using multiple linear regression models at the gene level, demonstrated Soil Diversity had significant effects (α=0.05) on the transcription of a broad set of CAZyme genes, in particular many glycoside hydrolases (7/21 genes), carbohydrate esterases (3/6 genes), glycosyl transferases (9/20 genes), and carbohydrate-binding modules (2/2 genes) (Figure 4). Similarly, urea addition had a significant effect on many CAZymes, especially glycoside hydrolases (8/21 genes), and there was a significant interaction effect for numerous glycoside hydrolase (4/21 genes) and glycosyl transferases (4/20 genes). At the CAZyme function level, High Diversity Urea (+) microcosms had the highest total expression of glycoside hydrolases, glycosyl transferases, and S-Layer homology genes compared to all other treatments. Transcription of carbohydrate-binding modules were highest in High Diversity soils, though urea addition to Low Diversity microcosms led to high variation in the total expression of carbohydrate-binding modules and no significant difference between Low Diversity Urea (+) to High Diversity soils. Transcription of auxiliary activities was highest in high diversity treatments and urea addition led to an increase in transcription within the high diversity treatment.

**Figure 4:**
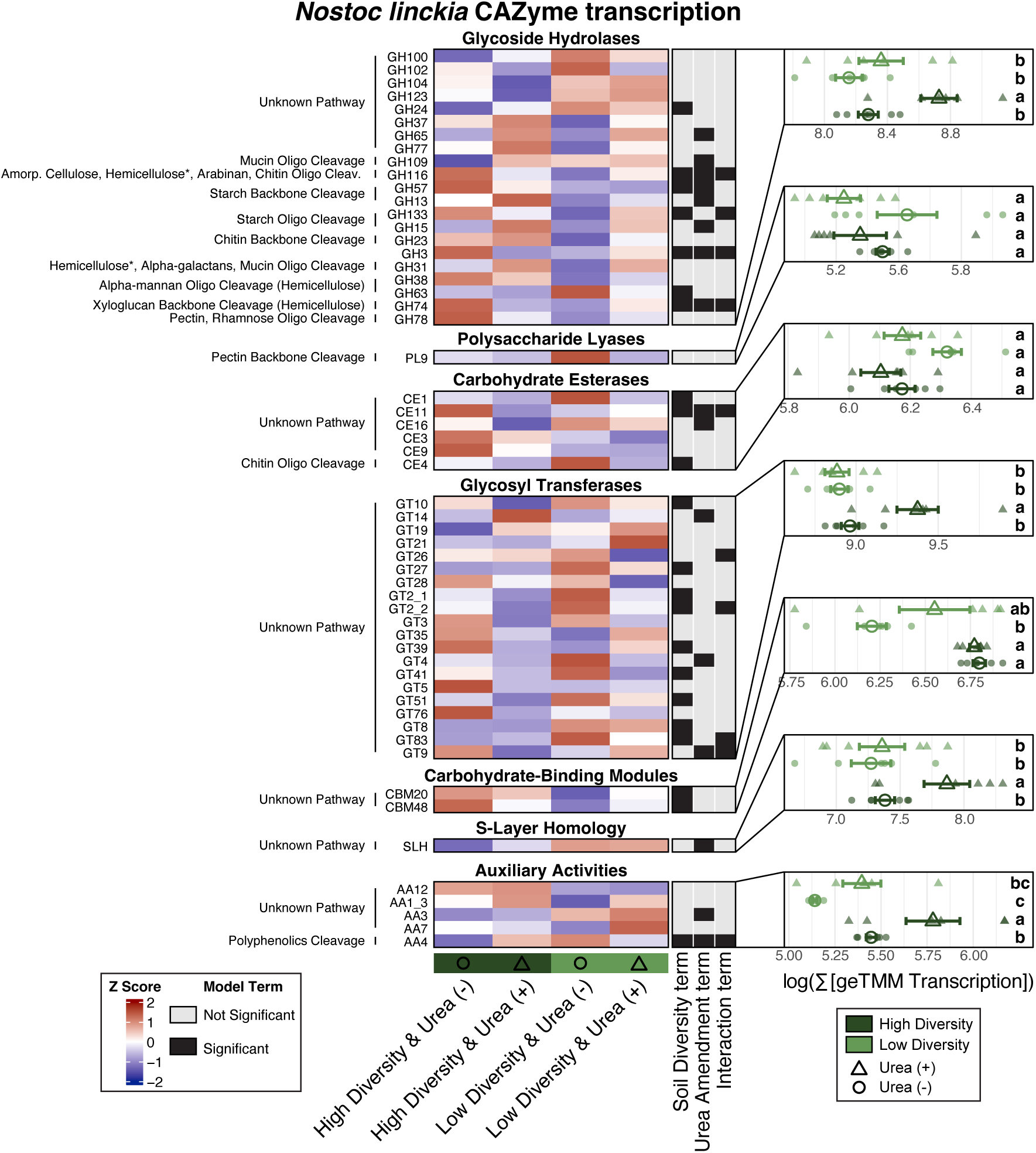
Heatmaps in left-most column represent z-scores of geTMM transcription values at the gene level (rows) for *N. linckia* CAZyme genes across the four treatments and faceted by pathways. Annotations at the gene level in middle column indicate significance (α=0.05, B-H corrected p-values) of model terms from multiple linear models across each gene. Right-most column shows the log of summed geTMM transcription values of genes among each pathway. Significance (α=0.05, B-H corrected p-values) between treatments were determined using ANOVA with Tukey-HSD post hoc tests and represented by grouping letters.

### Transcription of Nostoc linckia nitrogen metabolism and heterocyst formation genes

Because of the importance of DG1 and *N. linckia* on the nitrogen cycle in agroecosystems, we analyzed the transcription of nitrogen fixation, heterocyst-formation, and other nitrogen metabolism genes for *N. linckia*. At the gene level, Soil Diversity and Urea Amendment had a statistically significant effect (α=0.05) on only a few nitrogen fixation and heterocyst formation genes (Figure 5). However, at the pathway level, Urea Amendment had a significant effect on the transcription of the Mo- and V-nitrogenases where, across both diversity treatments, urea addition reduced expression of both nitrogenases. Further, V-nitrogenase expression displayed a soil diversity x urea amendment interaction effect, with no difference between Low Diversity Urea (+) and High Diversity Urea. Urea addition yielded lower transcription of HGL and PG remodeling genes in High Diversity microcosms only. No statistical difference in HEP layer genes was observed between either urea amendments or soil diversity levels, though High Diversity Urea (+) had statistically lower total transcription compared Low Diversity Urea (-). Additionally, no statistical differences were found between treatments for the total transcription of either heterocyst initiation and commitment or oxygen regulation genes.

**Figure 5:**
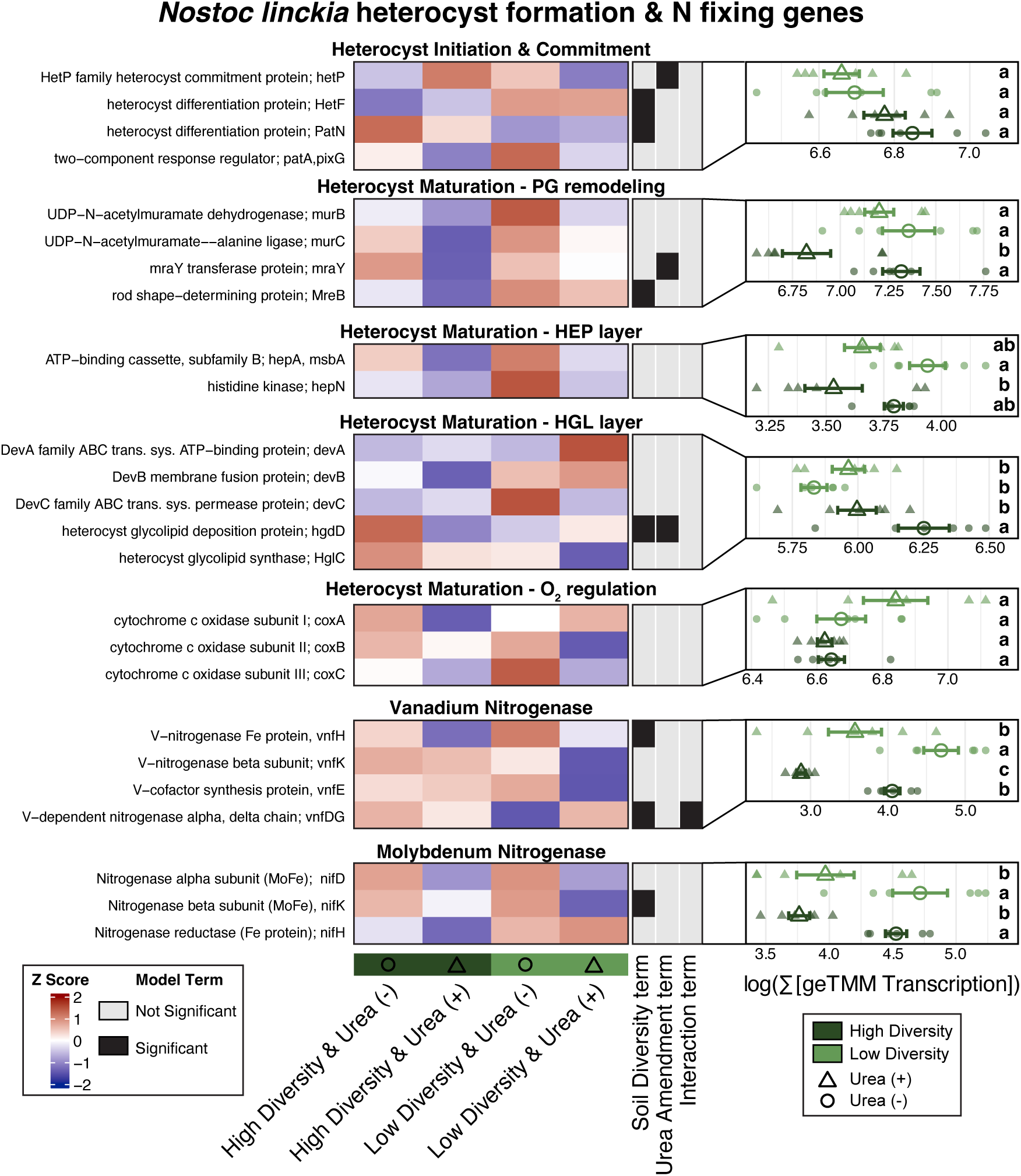
Heatmaps in left-most column represent z-scores of geTMM transcription values at the gene level (rows) for *N. linckia* heterocyst formation and nitrogen fixing genes across the four treatments and faceted by pathways. Annotations at the gene level in middle column indicate significance (α=0.05, B-H corrected p-values) of model terms from linear models across each gene. Right-most column shows the log of summed geTMM transcription values of genes among each pathway. Significance (α=0.05, B-H corrected p-values) between treatments were determined using ANOVA with Tukey-HSD post hoc tests and represented by grouping letters.

In addition to nitrogen-fixation and heterocyst formation genes, Soil Diversity and Urea Amendment drove significant differences in the transcription of other inorganic and organic nitrogen metabolism genes for *N. linckia* (Figure 6). At the gene-level, Soil Diversity explained differences (α=0.05) in the expression in many genes for the ornithine ammonium cycle, glutamate transformation, ammonium assimilation, assimilatory nitrate reduction, and nitrate/nitrite transport, while urea amendment had significant effects (α=0.05) on several genes within the ornithine ammonium cycle, cyanophycin degradation and synthesis, assimilatory nitrate reduction, and nitrate/nitrite transport.

**Figure 6:**
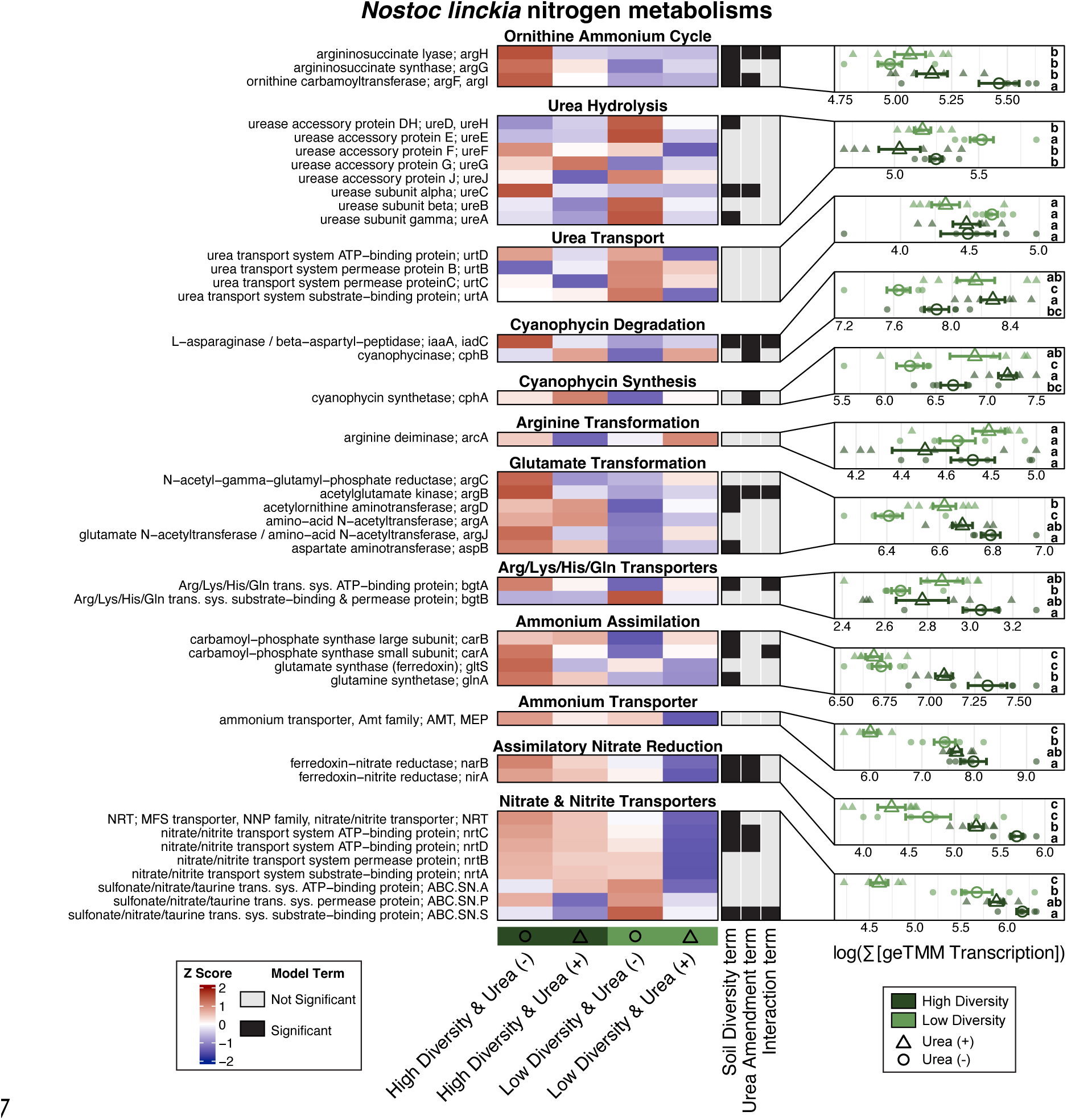
Heatmaps in left-most column represent z-scores of geTMM transcription values at the gene level (rows) for *N. linckia* nitrogen cycling genes across the four treatments and faceted by pathways. Annotations at the gene level in middle column indicate significance (α=0.05, B-H corrected p-values) of model terms from multiple linear models across each gene. Right-most column shows the log of summed geTMM transcription values of genes among each pathway. Significance (α=0.05, B-H corrected p-values) between treatments were determined using ANOVA with Tukey-HSD post hoc tests and represented by grouping letters.

Soil Diversity and Urea amendment had important effects on the total transcription of many organic nitrogen metabolisms at the pathway level. Specifically, *N. linckia* had a higher total expression of genes for the ornithine ammonium cycle in High Diversity Urea (-) compared to all other treatments. Low Diversity Urea (-) had higher total transcription of urea hydrolysis genes than all other treatments, though no treatment had a significant difference in total transcription of genes for urea transport. Similarly, no treatment differed in arginine transformation. In unfertilized soils, glutamate transformation genes were more highly expressed in High than Low Diversity microcosms; however, under urea fertilization, diversity treatments did not significantly differ. In low diversity microcosms, urea addition increased glutamate gene transcription compared to unfertilized samples. Total expression of Arg/Lys/His/Gln transporters was higher in high diversity microcosms compared to low diversity microcosms, however this difference was not observed with urea fertilization. Additionally, urea fertilization increased the total transcription of genes for both cyanophycin synthesis and degradation within High or Low Diversity treatments. Furthermore, High Diversity Urea (+) microcosms had higher total expression of cyanophycin synthesis and degradation that Low Diversity Urea (-).

In addition to organic nitrogen metabolism, the total transcription of inorganic nitrogen cycling genes by *N. linckia* were modulated by soil diversity and urea amendment. Here, High Diversity treatments had higher total transcription of genes for ammonium assimilation and assimilatory nitrate reduction compared to Low Diversity treatments, and the addition of urea lessened this effect in the High Diversity treatments. Similarly, the transcription of genes for ammonium and nitrate/nitrite transport were highest in High Diversity treatments, though Low Diversity Urea (-) was not statistically different than High Diversity Urea (+).

### Transcription of biofilm, extracellular polymeric substances, and hormogonium polysaccharide in Nostoc linckia

In order to determine how the experimental treatments may affect biofilm formation of *N. linckia*, a key potential function of DG1 as an inoculant, genes for production and secretion of extracellular polymeric substances (EPS) were explored (Figure 7). Additionally, one polysaccharide secretion gene, HpsB, was annotated to be specific for hormogonium polysaccharide secretion (HPS) and was included as *N. linckia* is capable of forming specialized motile cells called hormogonium. Soil Diversity had a significant effect on the transcription of HPS secretion, EPS, and polysaccharide transport genes, while the interaction of Soil Diversity and Urea Amendment had a significant effect on many of the same genes of EPS biosynthesis and polysaccharide transport. Low Diversity Urea (-) had higher total transcription of genes for EPS biosynthesis and polysaccharide transport than all other treatments. Similarly, Low Diversity Urea (-) had the highest total expression of HPS secretion and PEP-CTERM processing though there was no significant difference between urea amendments within Low Diversity treatments for HPS secretion or PEP-CTERM processing. Further, there was no significant difference between High Diversity Urea (-) and Low Diversity Urea (-) for PEP-CTERM processing.

**Figure 7:**
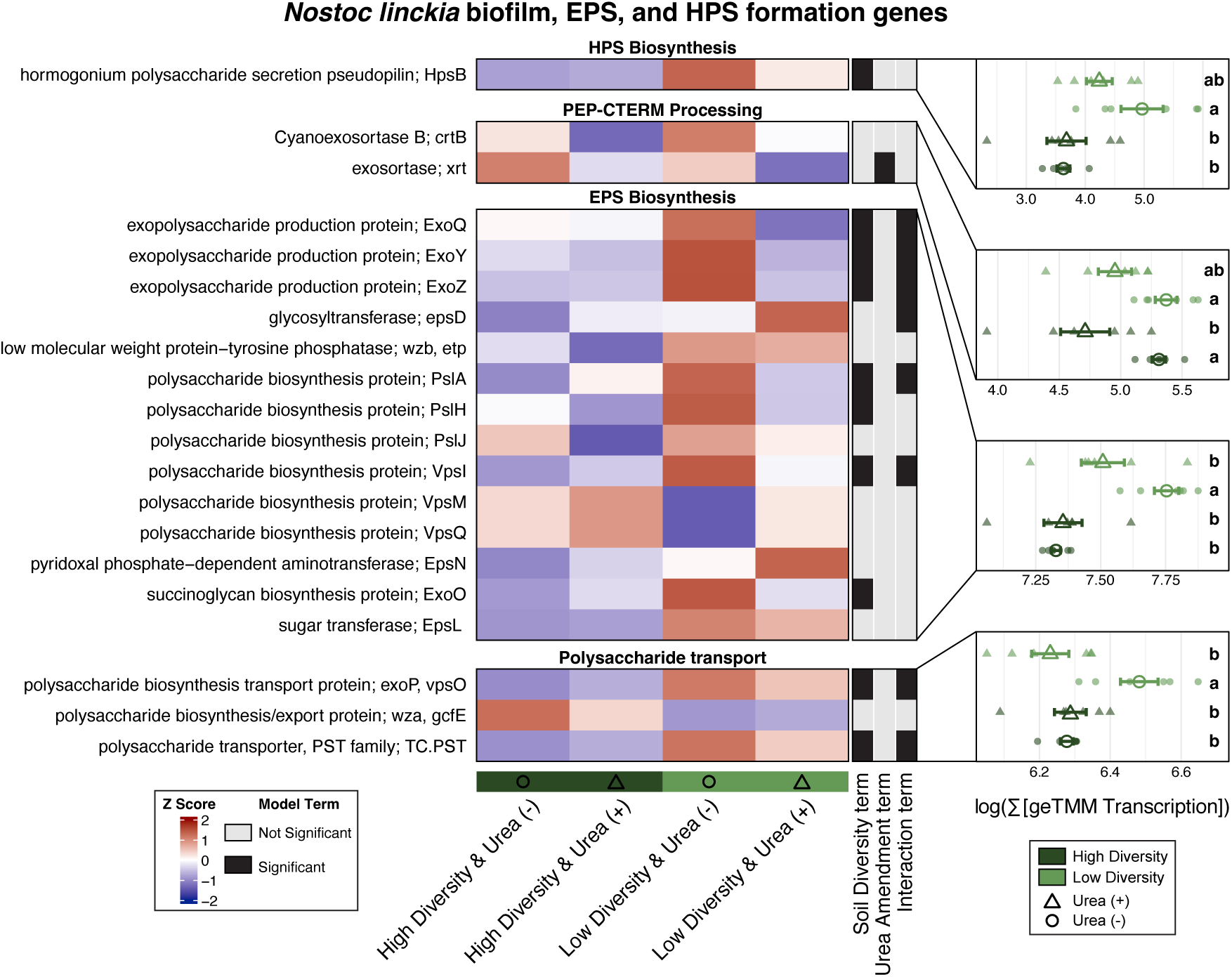
Heatmaps in left-most column represent z-scores of geTMM transcription values at the gene level (rows) for *N. linckia* EPS- and HPS-related genes across the four treatments and faceted by pathways. Annotations at the gene level in middle column indicate significance (α=0.05, B-H corrected p-values) of model terms from multiple linear models across each gene. Right-most column shows the log of summed geTMM transcription values of genes among each pathway. Significance (α=0.05, B-H corrected p-values) between treatments were determined using ANOVA with Tukey-HSD post hoc tests and represented by grouping letters.

## Discussion

The use of microbial inoculants for sustainable agriculture and restoration ecology is promising and has long been practiced; however, the establishment and functional performance of introduced microorganisms or microbiomes in the field can be unreliable, likely due to biotic and abiotic factors in recipient soils^9,52–55^. In this study, we tested whether resident soil microbiome diversity and urea fertilization history affected the establishment and transcription of a Cyanobacteria-dominated soil inoculant (DG1). We measured membership, relative abundance, and transcriptional activity of DG1 organisms in manipulated microcosms and assessed differences in the transcription of genes relevant for inoculant functioning (e.g. carbon-, nitrogen-cycling, CAZyme, and biofilm/EPS genes) of the dominant Cyanobacteria of DG1, *N. linckia*. Together, these results help clarify how soil biotic and abiotic factors influence inoculant establishment and functioning.

### Soil diversity and urea fertilization modulate the establishment and activity of DG1 members

In our microcosm experiment, resident soil diversity and urea fertilization history clearly influenced the ability of DG1 organisms to establish in recipient soils. Based on ASVs matched to DG1 organisms, four of the six heterotrophs in the DG1 inoculant failed to reliably establish in any of the experimental treatments. The two remaining DG1 heterotrophs, *Stenotrophomonas* sp. and *Ensifer* sp., established only in Low Diversity microcosms. Urea addition further modulated the ability of *Ensifer* sp. to establish in low diversity microcosms where it was only detected with added urea. In contrast to the heterotrophs of DG1, *N. linckia*, the dominant photosynthetic diazotroph present in DG1, was capable of establishing equally across all treatments regardless of soil diversity or urea addition. Additionally, these trends based on ASV data were further supported by transcript mapping data which demonstrated the proportion of genome transcribed and average gene transcription of each DG1 MAG is accordant with ASV detection in our microcosms.

Work on invasion repeatedly emphasizes that resident community biodiversity is a key driver of resistance to invasion^56–60^. Here the diversity-invasion relationship predicts the susceptibility of a community to invasion (i.e., invasion of an inoculant) is negatively related to the diversity of the resident community^61–63^. While at larger spatial scales this relationship can be neutral or even positive^64–68^, many recent studies on microbial invasion among varied ecosystems have reliably demonstrated a negative relationship between susceptibility to invasion and recipient community diversity^69^. The presence or absence of available niche space and resource availability within the recipient community for the invader is thought to drive this pattern^57,70–75^, where higher diversity communities exhibit higher niche complementarity and/or likelihood of a sampling effect^76^. Furthermore, the probability of invasion success is increasingly understood to depend on traits of both the invader and the recipient community^71,73,77–80^. In our study, the ability of inoculant members to establish in high diversity soils was trait-dependent, with clear contrasts between phototrophs (i.e., *N. linckia*) and the remaining DG1 heterotrophs. Microcosms in our study had little to no prior colonization of phototrophs before DG1 inoculation (Supplemental Figure 3), which, in addition to the capacity of DG1 to construct niche space through the production of a soil surface film, likely explains the reliability of *N. linckia* to colonize recipient soils regardless of soil diversity. Mawarda et al 2022^77^ demonstrated that some spore-forming bacterial inoculants can escape a negative diversity-invasion relationship by avoiding competition in the recipient community. Here, we build on this idea by proposing bacterial phototrophy as a trait that could yield a neutral diversity-invasion relationship when niche space for phototrophs is available. Conversely, we find that recipient soil diversity distinctly affected the colonization ability of *Stenotrophomonas* sp. and *Ensifer* sp. which are heterotrophic organisms. Consistent with the diversity-invasion relationship, high diversity microcosms were likely to harbor a more diverse set of heterotrophs than low diversity soils, and thus, less niche space available for DG1 heterotrophs.

### Soil diversity and urea fertilization alters the genome-wide expression of DG1

Following successful establishment in recipient soils, an inoculant is expected to induce a desired outcome by exhibiting specific phenotypes and performing targeted metabolisms. Reliability of successful inoculant performance is clearly context-dependent, where effective functioning is contingent on the soil biotic and abiotic conditions. Only a limited number of studies have tracked the shifts in gene transcription of established inoculants^80–82^, which have demonstrated that gene expression of established inoculants varies between liquid culture and soil environments^82^ and across different soil types^80,81^. We find that soil diversity explains a larger proportion of the variation in genome-wide gene expression of the DG1 inoculant compared to urea fertilization history (Figure 1), indicating the recipient soil community has a large effect on the overall metabolism of inoculants following successful establishment. This pattern likely reflects the wide range of potential interactions that inoculated microbes encounter across different soil microbiomes^80,83^, which can lead to different metabolic responses by the inoculant^81^. Here, among consistent soil abiotic conditions, our work highlights resident soil diversity as a key lever that modulates genome-wide expression patterns, even after successful establishment.

Nevertheless, soil abiotic properties are commonly cited as key determinants of inoculant effectiveness^84,85^. In our study, although the effects are smaller than soil diversity, urea fertilization and the interaction of soil diversity and urea fertilization each explain a modest portion of the variation in overall DG1 inoculant transcription (Figure 1). It remains unclear if these patterns would hold true for inoculants that exhibit different traits than that of the DG1 organisms or under different abiotic pressures. It is feasible that abiotic pressures other than urea fertilization might have a larger effect on the overall DG1 transcription, as DG1 is dominated by a diazotrophic Cyanobacteria that can acquire fixed nitrogen and carbon, and thus potentially less limited by added nitrogen. As such, future work should investigate the effect of soil diversity on genome-wide expression across inoculants with different metabolic traits and different abiotic conditions.

### Soil diversity and nitrogen legacy jointly transforms N. linckia carbon and nitrogen metabolism

As a phototrophic diazotroph, *N. linkcia* has the potential as a prospective inoculant to add fixed carbon and nitrogen to agroecosystems and thus, we specifically explored the impact of resident soil diversity and urea fertilization on the transcription of carbon and nitrogen cycling genes. In the absence of added nitrogen (via urea), high diversity resident microbiomes push *N. linkcia* toward a more active phototrophic metabolism (Table 1). Total transcription of PSII, PSI, cytochrome b6f, carboxysome, RuBisCO, and photorespiration pathways were all increased in high diversity compared to low diversity treatments when urea was not added to microcosms. This suggests that potential mutualistic interactions with soil members in high diversity soils may play an important role in carbon fixation by *N. linckia*. Indeed, many Cyanobacteria, including *Nostoc* spp.^86^, have been shown to upregulate photosynthesis and carbon fixation genes when participating in mutualistic interactions with other microbes^87–89^. Interestingly, urea addition attenuated these soil diversity driven effects, resulting in no difference in many photosynthesis and carbon fixation pathways between diversity treatments. Photosynthesis by non-diazotrophic cyanobacteria can be limited by available nitrogen^90–92^ and, though a diazotroph, urea fertilization increased total transcription of PSI and cytochrome b6f by *N. linckia* in low diversity treatments. This potentially indicates a competitive release from competitors for urea uptake in the diverse soil microbiome; however, we found no increase in transcription of urea transport or hydrolysis in the urea amended low diversity microcosms.

**Table 1:** Conceptual summarization of the relative effects of treatments on key carbon and nitrogen metabolism pathways for *N. linckia*.

| **Function** | **Low Diversity**<br>**Urea (-)** | **Low Diversity**<br>**Urea (+)** | **High Diversity**<br>**Urea (-)** | **High Diversity**<br>**Urea (+)** |
| --- | --- | --- | --- | --- |
| **Phototrophy** | Low | Medium | High | High |
| **CAZymes** | Low | Medium | Low | High |
| **Nitrogenase** | High | Low | High | Low |
| **Inorganic N**<br>**assimilation** | Low | Low | Medium | High |
| **Glutamate transformation** | Low | Medium | Medium | High |
| **Ornithine ammonium cycle** | Low | Low | Low | High |
| **Cyanophycin** | Low | Medium | Medium | High |

In order to test whether soil diversity and urea fertilization alter organic carbon metabolisms in *N. linckia*, we further assessed the transcription of CAZyme genes. High diversity microcosms with nitrogen added had the highest total expression of genes for glycoside hydrolases, glycosyl transferases, carbohydrate binding modules, and S-layer homology, and auxiliary CAZyme activities (Table 1). This is consistent with prior work that has shown nitrogen addition to soil microbial communities frequently stimulates the expression of CAZymes^93–95^. However, urea fertilization did not increase the total transcription of any CAZyme family in low diversity treatments, suggesting the effect of nitrogen addition on *N. linckia* carbohydrate metabolism depended on the surrounding microbial community. In particular, the elevated transcription of glycoside hydrolases and glycosyl transferases in high diversity, urea amended soils suggests increased carbohydrate degradation and synthesis/modification under these conditions. Because the DG1 inoculant forms a biofilm on the soil surface, these shifts could reflect changes in extracellular polysaccharide dynamics. However, the total expression of *N. linckia* EPS biosynthesis and polysaccharide transport genes was not significantly higher in high-diversity, urea amended soils, suggesting that increased CAZyme transcription may instead reflect EPS remodeling, carbohydrate turnover, or interactions with soil organic matter rather than increased EPS production alone.

Beyond their broad effects on *N. linckia* carbon cycling, soil diversity and urea amendment also significantly shaped nitrogen cycling metabolisms. Urea fertilization resulted in lower expression of both Mo- and V-nitrogenases within each soil diversity level (Table 1). This agrees with the numerous work that has shown decreased nitrogenase transcription and activity following nitrogen addition to soils^96–99^. Further, the total transcription of genes for PG remodeling and HGL deposition were downregulated in urea amended microcosms, but only in high diversity treatments. This points to a case where in low diversity microcosms, expression of heterocyst forming genes was similar with or without added urea, but in amended soils nitrogenase expression was downregulated. A decoupling in heterocyst frequency and nitrogenase expression has been observed previously^100,101^ which may be the case in urea amended low diversity soils, though further work would be necessary to confirm this. Additionally, within soil diversity levels, the total transcription of cyanophycin synthesis and degradation genes was consistently higher in urea fertilized soils (Table 1). As cyanophycin is a nitrogen and carbon storage polymer often localized as granules at the poles of heterocysts, this indicates an increased flux of carbon and nitrogen through cyanophycin in urea fertilized soils.

While nitrogen fixation and heterocyst genes responded largely to urea fertilization, downstream nitrogen assimilation and storage pathways responded to both soil diversity and urea addition. In unfertilized microcosms, high diversity soils led to increased transcription of organic nitrogen cycling pathways (ornithine ammonium cycle, glutamate transformation) and inorganic nitrogen assimilation (assimilatory nitrate reduction and ammonium assimilation via GS/GOGAT and carbamoyl-P synthase) by *N. linckia* (Table 1). This may be attributed to a higher capacity of the diverse soil microbiome to cycle nitrogen, resulting higher soil nitrogen mineralization and nitrification; and, thus, production of inorganic nitrogen which *N. linckia* assimilates. Further studies should quantify inorganic and organic nitrogen concentrations in relation to DG1 and *N. linckia* inoculation on low and high diversity soil microbiomes or with microbiomes with varying activity of ammonification and nitrification. Urea addition to high diversity soils lessened the effect of increased inorganic nitrogen assimilation pathway expression, though these pathways remained more highly expressed than in low diversity soils (Table 1). Notably, Patra et. al. (2006)^102^ suggests nitrogen fixation is promoted in bulk soils with low microbial activity and low net nitrogen mineralization. This is consistent with our unfertilized low diversity treatment, both total nitrogenase expression and inorganic nitrogen assimilation were lowest in unamended low diversity soils, potentially owing to lower nitrogen mineralization as a product of low soil diversity. These patterns highlight a potential increased reliance on nitrogen fixation as opposed to inorganic nitrogen assimilation in unfertilized low diversity soils by *N. linckia*.

### Implications for the deployment and use of soil inoculants

We find that resident soil diversity and urea fertilization interact to induce important shifts in key carbon and nitrogen metabolisms by *N. linckia* that are relevant to inoculant functioning of the DG1 consortium. Such variation in gene transcription could potentially explain, in part, the variability in inoculant performance observed in the field. This microbiome and nutrient status specific effect on functioning may complicate the development of tailored, “field-specific” inoculants which have been proposed as a solution to increase inoculant reliability^103^. Furthermore, this work highlights the importance of considering the set of dominant traits of a potential inoculant and how these traits might interface with the resident soil ecology (i.e., niche occupancy) to impact establishment and functioning *in situ*. For example, we find that the establishment of the dominant diazotrophic phototroph in DG1 is not affected by soil diversity or urea fertilization, potentially because of the unique niche space it occupies in the soil community. On the other hand, soil diversity impacts the establishment of the minor abundance heterotrophs possibly due to increased competition with other soil heterotrophs.

## Conclusions

By leveraging genome-resolved metatranscriptiomics, our study uniquely provides direct evidence that the resident soil microbiome and urea fertilization can induce broad shifts in gene transcription patterns of a potential inoculant and interact to reprogram the expression of crucial carbon and nitrogen metabolism pathways. We find that when interacting with a complex community, inoculant phenotypes are not fixed and are affected by soil biology and abiotic conditions. This suggests that successful establishment alone may be insufficient to predict inoculant performance, because the functional traits expressed by an inoculant may depend strongly on the soil environment into which it is introduced. As a result, the benefits of inoculation are likely to emerge from interactions between the inoculant, the resident microbiome, and local soil physicochemical conditions. In all, future work on inoculant development should consider the ecological interactions (both biotic and abiotic) a prospective inoculant might experience in the field and how these interactions might affect establishment and functioning of the inoculant.

## Abbreviations

DG1: Dark Green 1;
(SSC): Soil Surface Consortia;
RIN: RNA Integrity Number;
US-DOE: United States Department of Energy;
JGI: Joint Genome Institute;
MAG: Metagenome-assembled Genome;
ASV: Amplicon Sequence Variant

## Acknowledgements

This work was supported by USDA NIFA Hatch Projects 1016233 and 1003346, as well as a Pennsylvania State University CAS RAIN grant. Metatranscriptomic sequencing was enabled by U.S. Department of Energy Joint Genome Institute CSP 503310. We thank Roxanne Lease for help in setting up incubation infrastructure and Timothy Peoples for integral assistance in performing DNA extractions.

