## Supplementary Information for "Resident soil microbial diversity and urea amendment legacy interact to shape the composition and expression of a surface film–forming soil inoculant"

**Supplementary Material**

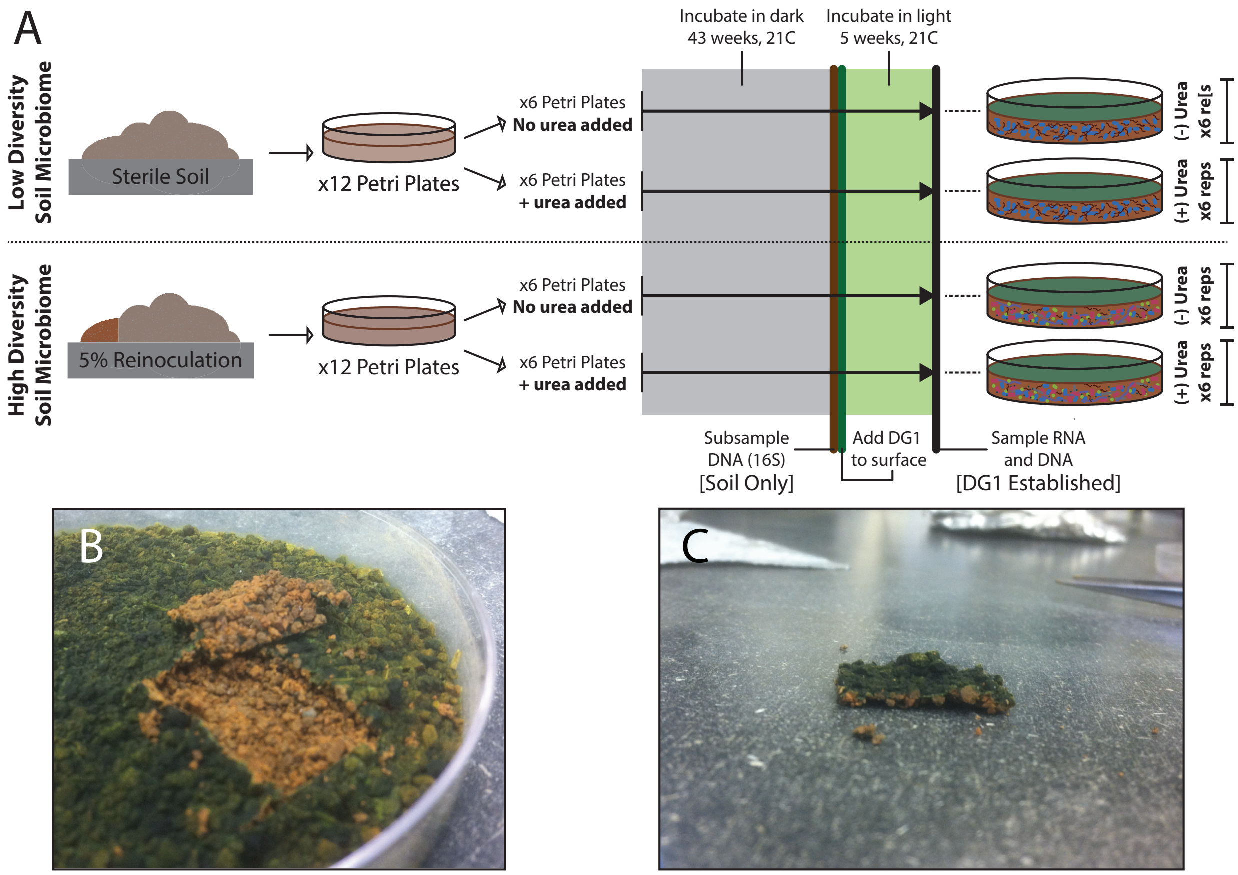


**Supplemental Figure 1: A)** Schematic showing how microcosms were prepared. Both High and Low Diversity soils began with the same batch of twice autoclaved soil. Soil for High Diversity was reinoculated with the same non-autoclaved soil at 5% w/w to reintroduce a diverse soil microbiome. Soils were distributed to petri plates and half received a urea amendment. Prepared microcosms were incubated in the dark for 43 weeks to reestablish microbiomes and then subsampled for DNA extractions (Sampling Phase, Soil Only). DG1 was pipetted to microcosm surfaces and allowed to establish for 5 weeks under light before destructively harvesting for DNA and RNA extractions (Sampling Phase, DG1 Established). **B)** Photo showing a microcosm patch to be used for DNA and RNA extractions (Sampling Phase, DG1 Established). **C)** Example of a sampled patch from a test microcosm. The DG1 inoculant grows as a biofilm on the surface of the soil and embeds some attached soil particles.

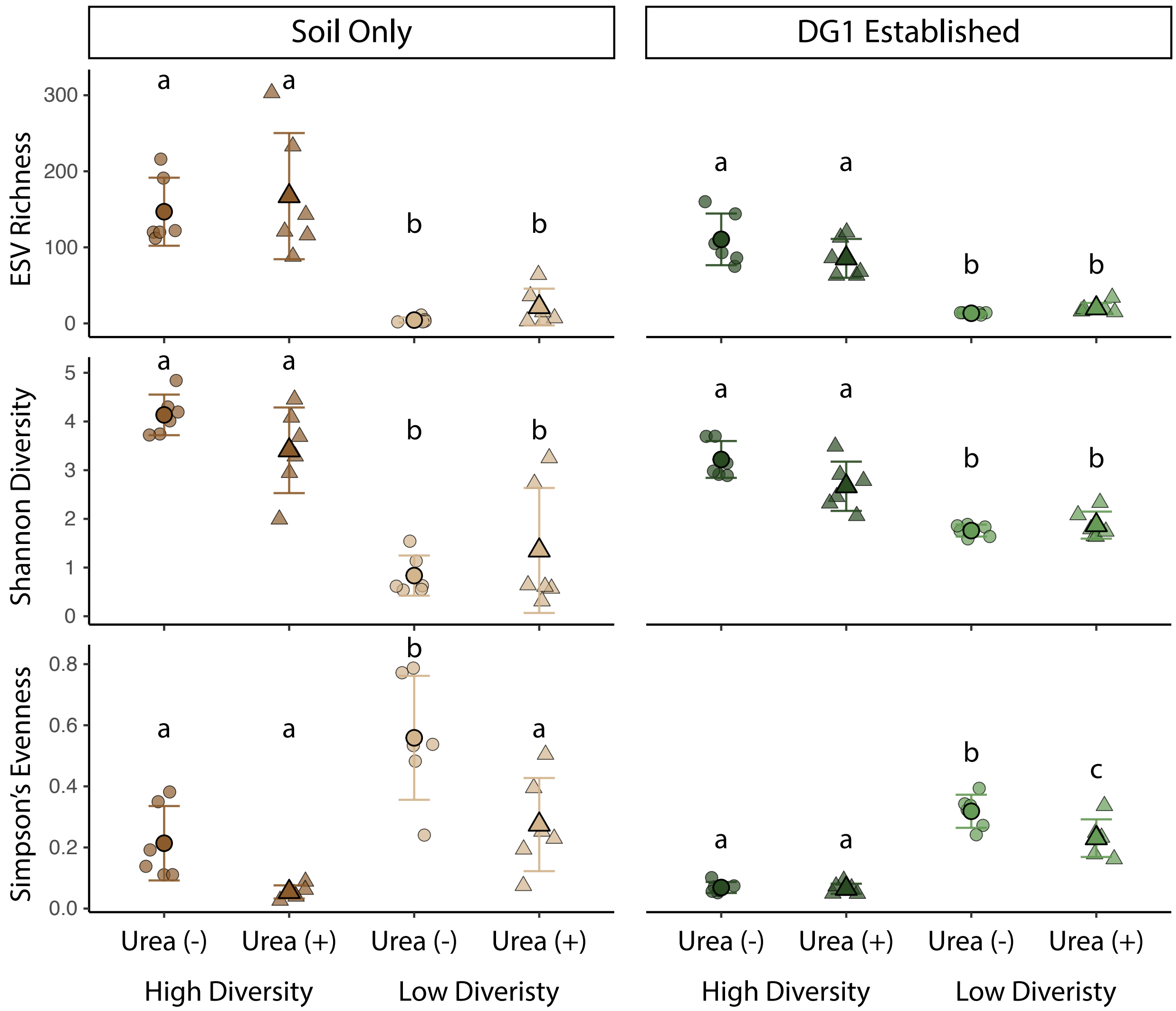


**Supplemental Figure 2**: Alpha diversity metrics for microcosms prior to DG1 addition (Left panel) and at the end of the experiment after DG1 addition (Right panel). Brown (High Soil Diversity) and tan (Low Soil Diversity) points indicate microcosms prior to DG1 addition, while microcosms at the end of the experiment with DG1 added are displayed in Dark Green (High Soil Diversity) and Light Green (Low Soil Diversity). Triangles represent microcosms that received urea amendment, while circles show microcosms with no added urea. ANOVA with Tukey-HSD post-hoc tests were performed within each alpha diversity metric and sampling phase to test differences between treatments as indicated by grouping letters. Omnibus P-values were adjusted via Benjamini-Hochberg and significant considered at α=0.05.

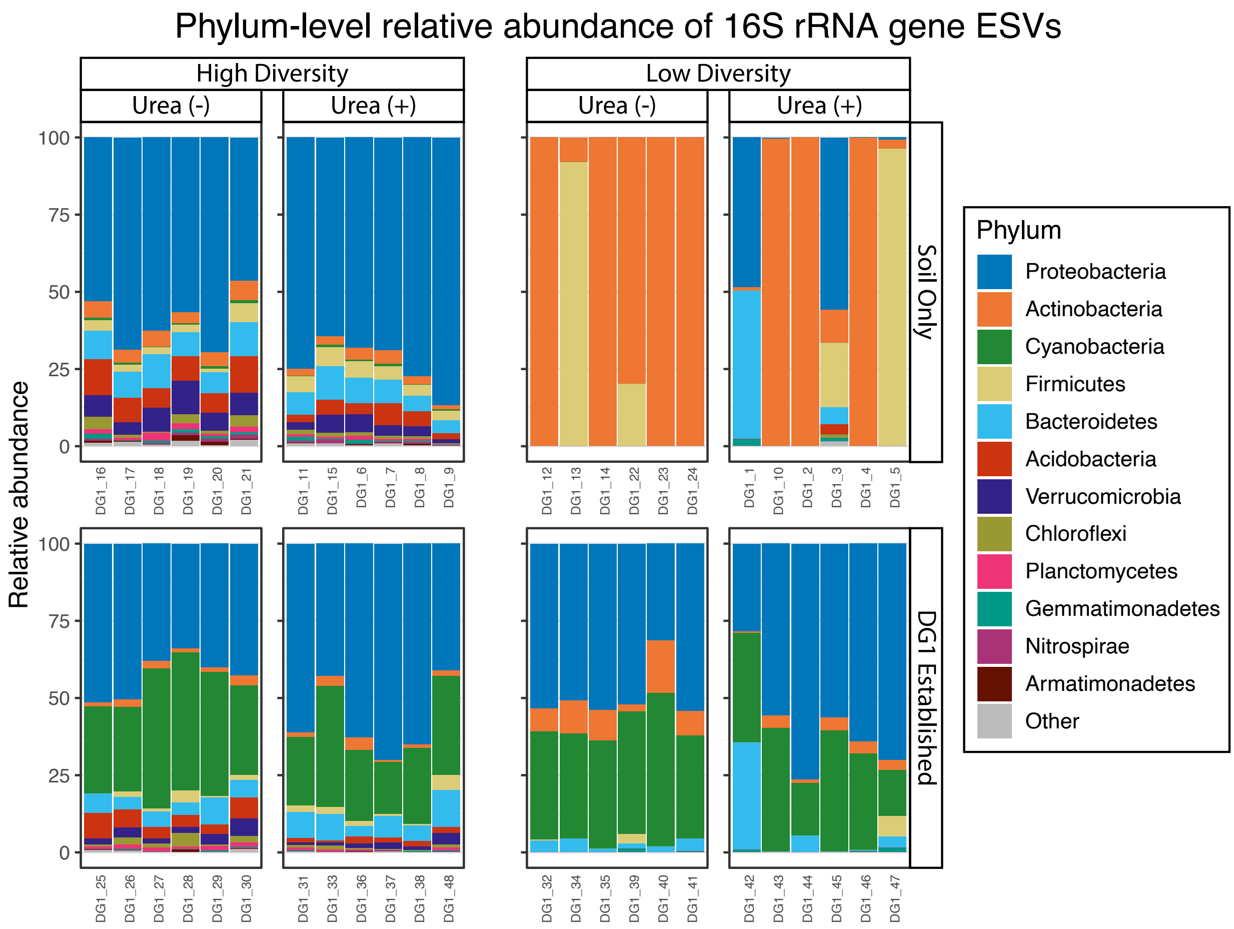


**Supplemental Figure 3**: Relative abundance plots of ASVs at the Phylum level. Plots are faceted by Soil Diversity and Sampling Phase. The top 12 most abundant phyla were plotted and the remaining ASVs were aggregated to “Other.”

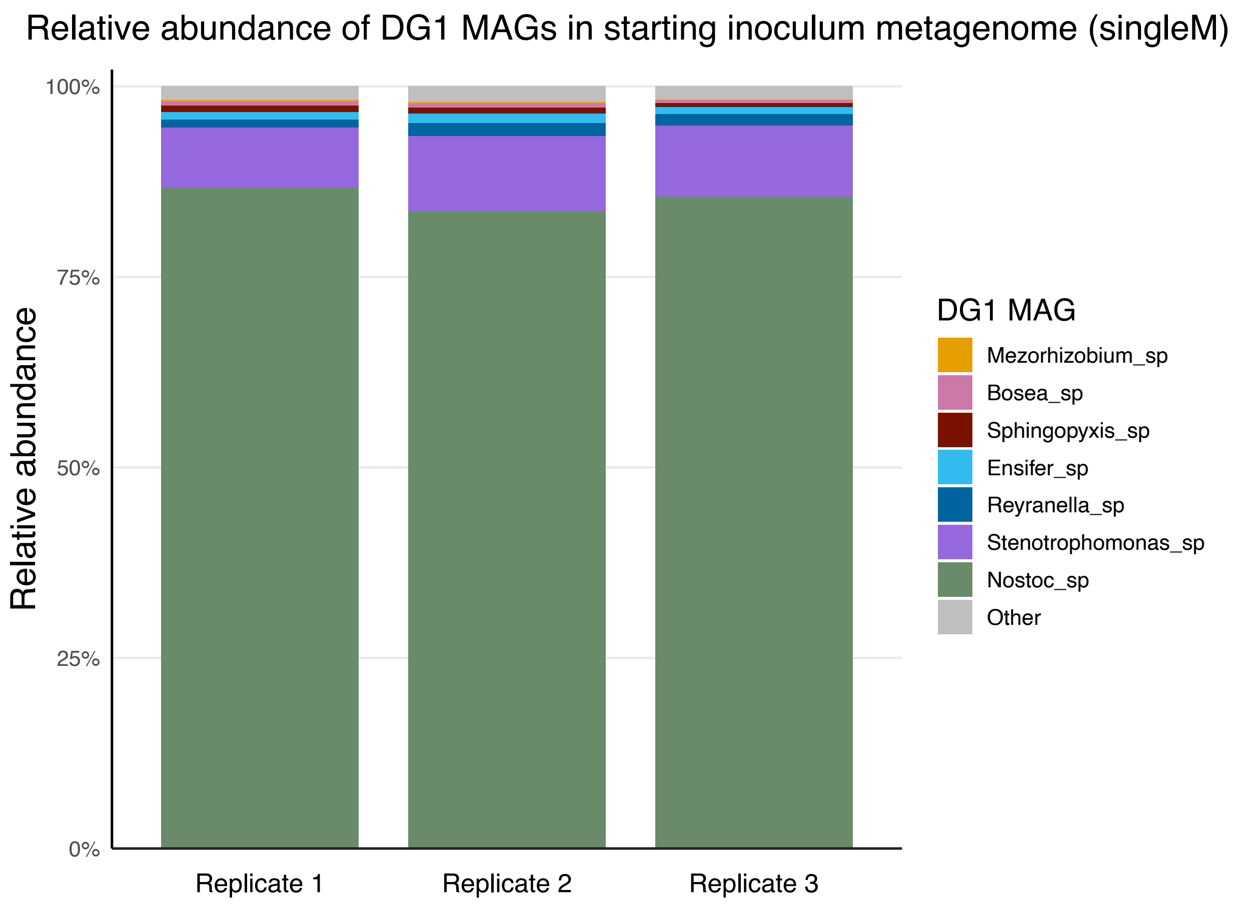


**Supplemental Figure 4**: Relative abundance of DG1 MAGs in technical replicates of the DG1 inoculant metagenomes from singleM analysis. “Other” denotes the aggregated sum of abundance of any bacterial taxa not included in the seven MAGs recovered from the metagenomes.

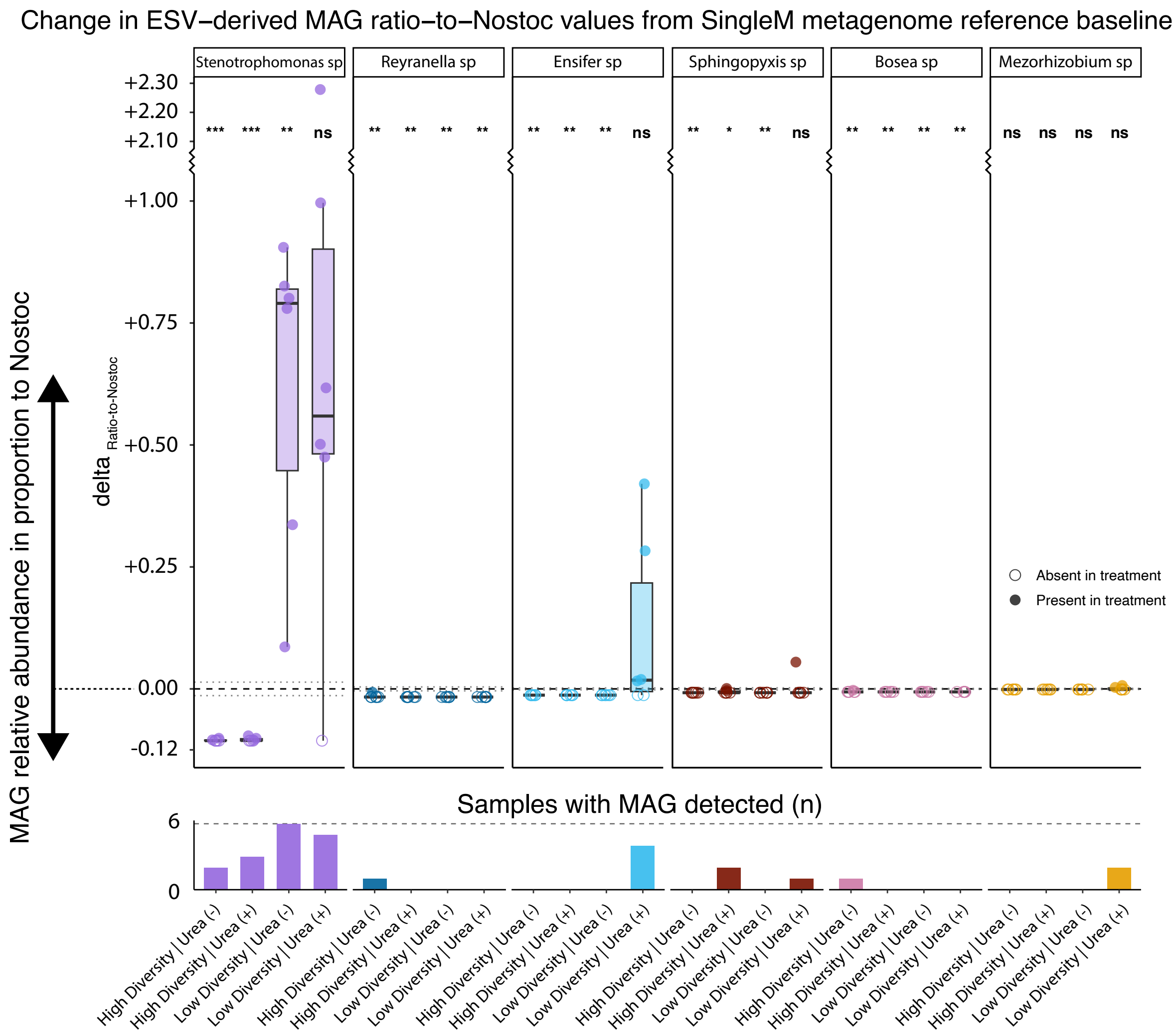


**Supplemental Figure 5**: Top: Change in MAG ratio-to-Nostoc values at time of sampling (points and boxplots) compared to MAG ratio-to-Nostoc in the starting inoculum (black dashed line). Ratios at time of sampling were calculated using ASV-derived relative abundance of DG1 MAGs and ratios from the starting inoculum were estimated using singleM. Closed circles indicate MAG was detected in the microcosm replicate at the time of sampling while open circles indicate the MAG was not detected. Bottom: Bar plots of the frequency in which the MAG was detected among the 6 microcosm replicates per treatment. All data are faceted by DG1 MAG.

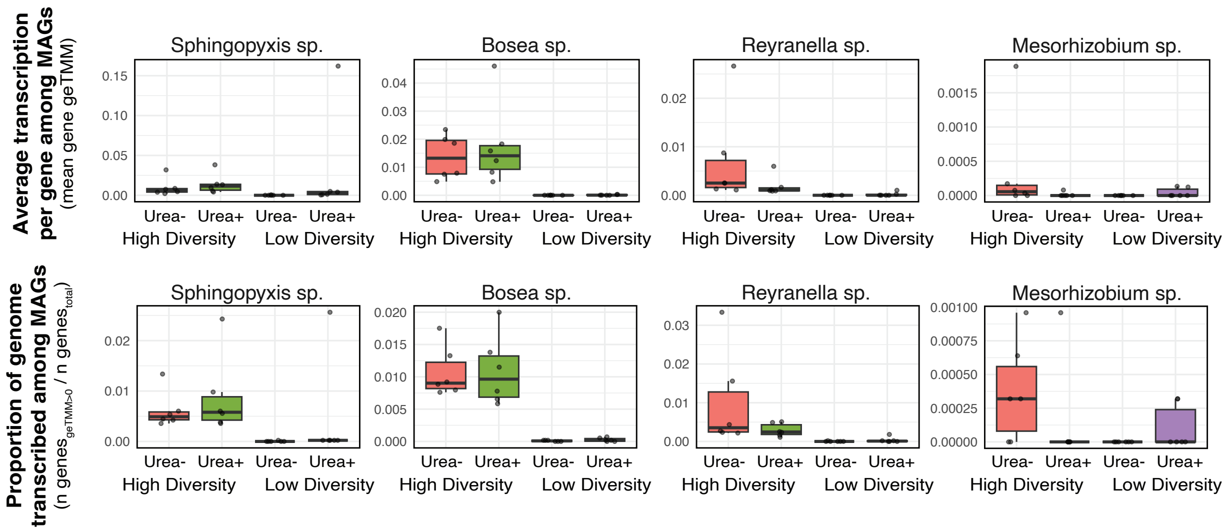


**Supplemental Figure 6**: Top: Boxplots of the average gene transcription (geTMM) among the minor abundance DG1 MAGs. Bottom: Boxplots of the proportion of the genome with mapped transcripts assigned among the minor abundance DG1 MAGs.

**Supplemental Table 1**: Summary information regarding MAGs.

| MAG | Completeness (%) | Contamination (%) | 23S rRNA count | 5S rRNA count | 16S rRNA count | tRNA count | Assembly Length (bp) | Contigs (n) | Largest Contig (bp) | N50 | GC (%) | Pred. Genes (n) |
|---|---|---|---|---|---|---|---|---|---|---|---|---|
| *Sphingopyxis* sp. | 100 | 2.1 | 1 | 1 | 0 | 47 | 4684941 | 140 | 141099 | 47720 | 63.81 | 4488 |
| *Bosea* sp. | 99.87 | 0.1 | 1 | 1 | 0 | 45 | 5626147 | 313 | 106260 | 26780 | 67.16 | 5653 |
| *Reyranella* sp. | 99.64 | 0.41 | 1 | 1 | 1 | 44 | 5623802 | 27 | 723106 | 287762 | 66.03 | 5516 |
| *Nostoc linckia* | 99.98 | 1.35 | 1 | 0 | 0 | 74 | 7771479 | 131 | 276682 | 103539 | 40.63 | 6429 |
| *Ensifer* sp. | 94.95 | 1.41 | 1 | 1 | 1 | 46 | 6527224 | 41 | 533427 | 256430 | 61.96 | 6050 |
| *Stenotrophomonas* sp. | 100 | 0.36 | 1 | 1 | 1 | 68 | 5057699 | 21 | 635102 | 394614 | 66.35 | 4587 |
| *Mesorhizobium* sp. | 41.11 | 0.1 | 0 | 0 | 0 | 19 | 2704125 | 662 | 13212 | 4175 | 65.76 | 3130 |

**Supplemental Table 2**: Summary statistics of MAG relative abundance in DG1 inoculant metagenome technical replicates (n=3) from singleM.

| MAG | hits mean | hits sd | hits mean relative abundance | hits mean relative abundance sd |
|---|---|---|---|---|
| *Mezorhizobium* sp. | 15.33 | 12.58 | 0.11 | 0.09 |
| *Bosea* sp. | 73.33 | 11.68 | 0.53 | 0.07 |
| *Sphingopyxis* sp. | 91.33 | 23.46 | 0.67 | 0.18 |
| *Ensifer* sp. | 151.33 | 33.08 | 1.09 | 0.14 |
| *Reyranella* sp. | 199.00 | 62.39 | 1.42 | 0.32 |
| *Stenotrophomonas* sp. | 1256.67 | 262.73 | 9.04 | 1.03 |
| *Nostoc* sp. | 11749.33 | 988.40 | 85.26 | 1.54 |
| Other | 260.00 | 47.62 | 1.88 | 0.18 |
